# Spontaneous helping in pigs is mediated by helper’s social attention and distress signals of individuals in need

**DOI:** 10.1101/2023.03.17.533160

**Authors:** Liza R. Moscovice, Anja Eggert, Christian Manteuffel, Jean-Loup Rault

**Affiliations:** Institute of Behavioural Physiology, Research Institute for Farm Animal Biology (FBN), Dummerstorf, Germany; Institute of Genetics and Biometry, Research Institute for Farm Animal Biology (FBN), Dummerstorf, Germany; Institute of Animal Welfare Science, University of Veterinary Medicine, Vienna, Austria

**Keywords:** *Sus scrofa*, prosocial, local enhancement, cortisol, rescue, empathy

## Abstract

Helping behaviour is of special interest for prosociality because it appears to be motivated by the needs of others. We developed a novel paradigm to investigate helping in pigs (*Sus scrofa domesticus*) and tested 75 individuals in eight groups in their home pens. Two identical compartments were attached to the pen, equipped with a window, and a door that could be opened from the outside by lifting a handle. Pigs in all groups spontaneously opened doors during a five-day familiarization. During testing, each pig was isolated once from its group and placed in one of the two compartments, in a counter-balanced order. In 85% of cases, pigs released a trapped group member from the test compartment within 20 minutes (median latency = 2.2 minutes). Pigs were more likely and quicker to open a door to free the trapped pig than to open a door to an empty compartment. Pigs who spent more time looking at the window of the test compartment were more likely to help. Distress signals by the trapped pig increased its probability of being helped. Responses are consistent with several criteria for identifying targeted helping, but results can also be explained by selfish motivations.

## 1. Background

Helping behaviour, in which an individual assists another to reach an otherwise unachievable goal, is of special interest for the study of prosociality. Helping can be costly (1), and often does not provide immediate benefits to the helper, but rather appears to be motivated by an understanding of the needs of others, also referred to as targeted helping (2). Examples of helping are wide-spread across animal taxa in the wild, and include intervening on behalf of group members against predators (ants: (3); primates: (4); whales: (5)), releasing individuals caught in snares, traps or sticky seeds (ants: (6); wild boars (7); birds (1)) and defending group members from conspecific attacks (elephants: (8); primates: (9)).

Animal helping is also the subject of experimental research to identify common biological mechanisms underlying empathy, broadly defined as the ability to recognize the emotional states of others and generate appropriate responses (2,10). Animal helping and human empathic responses share several trait features (10), including that both are mediated by the processing of social affective cues in similar brain regions (rodents: (11), humans: (12)). Long-standing criticisms maintain that the available evidence is not sufficient to support empathy-like motivations underlying animal helping, and that more parsimonious explanations are possible (13–16). One major point of contention is about whether helping is truly goal-oriented, which would require showing not only that animals help in response to individuals in need, but also that observers modify helping appropriately when the target’s needs change or when they no longer require help (17). Other criticisms highlight issues with the design of helping experiments themselves, which can make it difficult to distinguish between prosocial and selfish motivations for helping, for example when helping provides access to desirable resources such as social partners (e.g. social reinforcement, 14,18), or novel locations (19-20), in an otherwise barren test environment. Moreover, some studies have assumed that their paradigms elicit distress without measuring responses of target individuals (e.g. 19), leading others to question whether these individuals are even in need of help (14,16,18). Interestingly, there is also evidence that high levels of distress by individuals in need may sometimes hinder helping (21-22). More studies are thus needed to disentangle the different motivations for helping behaviour (16).

We developed a novel helping paradigm that gives potential helpers and individuals in need more behavioural flexibility to express a range of responses, in an attempt to address some of these criticisms. Our paradigm is inspired by the work of Ben-Ami Bartal and colleagues (19), in which rats could open a door to release a partner trapped inside a restraint tube. However, our approach differs in several important aspects. Firstly, subjects are tested in social groups in their home environment (e.g. (23)), which improves ecological validity by giving animals the choice about whether, when and whom to help, while allowing for competing motivations (e.g. exploring, feeding, resting, social interactions with other group members). Secondly, subjects can simultaneously perform the same behaviour in a range of helping as well as non-helping contexts, which allows us to assess different motivations for door-opening. Finally, the trapped individual is able to move around and explore and can also maintain visual contact and interact with potential helpers through a mesh window, allowing us to investigate how different behavioural responses of trapped individuals influence helping (24).

We tested the paradigm in pigs, which are a promising model species for studying the mechanisms mediating helping behaviour. Pigs show evidence of emotional contagion when observing group members in stressful situations (25), but without opportunities to assist them. In addition, a case of spontaneous rescue behaviour was recently documented in wild boars (*Sus Scrofa*), the ancestors of domestic pigs (7). However, helping has not been systematically studied in pigs.

Our objectives were to determine whether pigs would spontaneously open a door to free a trapped group member, to identify the factors that influence door-opening behaviour, and to test whether pigs’ responses were consistent with targeted helping or selfish motivations. We followed the criteria suggested by Pérez-Manrique and Gomila (26) for demonstrating targeted helping in animals: i) Helpers should show an other-oriented reaction, including cognitive appraisal of the situation and a moderate level of arousal, consistent with emotional regulation; ii) Helpers should show a flexible helping response that is appropriate to the situation and iii) Helping should lead to an improvement in the situation for the individual in need.

We predicted that pigs would selectively and consistently open doors to release trapped group members, but not to access empty compartments. We further predicted that potential helpers would be more likely and quicker to help individuals: i) if they were more proficient in the novel door-opening behaviour, ii) if they were more attentive to the trapped individual and iii) if the trapped individual expressed greater need to be released, either by producing distress signals or by attempting to make social contact. We measured changes in salivary cortisol (sCORT) in trapped pigs, to test whether specific behavioural responses to being trapped were associated with greater activation of the hypothalamic-pituitary-adrenal axis, consistent with a physical or psychological stress response (reviewed in (27). Whenever the identity of helpers could be quickly confirmed without requiring video analysis, we also measured their sCORT concentrations soon after helping, to test whether helpers showed evidence for increases in cortisol, consistent with emotional contagion of distress from trapped pigs, or for more moderate levels of arousal, consistent with emotional regulation that can facilitate targeted helping (21,26).

## 2. Methods

### 2.1 Animals and husbandry

All research procedures were approved under the German Animal Welfare Act (German Animal Protection Law, §8 TierSchG) by the Committee for Animal Use and Care of the Agricultural Department of Mecklenburg-Western Pomerania, Germany (permit LALLF 7221.3-1-036/20).

Subjects were German Landrace pigs born and housed at the Experimental Pig Facility at the Research Institute for Farm Animal Biology (FBN) in Dummerstorf, Germany. Pigs were tested in four temporally-separated cohorts, based on the five-week interval between the births of new litters. Each cohort consisted of two social groups, for a total of n = 78 subjects. At 28 days of age, subjects were weaned and mixed into new social groups, by randomly selecting 3-5 pigs from each of two to three different litters to form groups of 9-10 individuals. Groups were sex-balanced and contained equal numbers of littermates from each contributing litter, such that each pig had either two (22% of group members) or four (44% of group members) relatives in their group. Groups within a cohort were housed in identical, neighbouring 2.8 × 2.8 m pens in the same room, with partial slatted flooring and a sleeping mat. Throughout the study, pigs in neighbouring pens did not have direct visual or physical contact but could exchange auditory and olfactory cues. Pigs were regularly given back numbers using livestock spray paint to facilitate individual identification. The room was equipped with multiple video cameras in order to cover the entire home pen and two additional compartments that were attached to the home pen for the experiment (see section 2.2). Pigs were tested between 46 – 55 days of age. Prior to testing, pigs were habituated to saliva sampling and had participated in one social play experiment within their social group (28). Of the 78 subjects, one died of unknown causes, and two subjects from the same group were removed from the study to be treated for illnesses before testing began. We report results for the remaining n = 75 subjects who completed the study.

### 2.2 Familiarization, separation and testing

Two weeks after weaning, pigs went through familiarization, with the goal to expose pigs to novel compartments and a novel door-opening mechanism. Each group within a cohort had a one-hour familiarization session per day, with groups tested consecutively in a counter-balanced order across five days. On each familiarization day, pigs were presented as a group in their home pen with two identical compartments (90 × 90 × 80 cm in height) that were temporarily attached to opposite sides of the home pen (**Fig. S1**). Compartments were open on top to facilitate placement of trapped pigs and video recording during testing. Each compartment had a mesh window and a door with a metal handle, placed at 20 cm height (**Fig. S2**). Opening the door required the use of 1.3 kg of force to lift the handle with the snout high enough to release an inner latch, causing the door of the compartment to swing open into the home pen. This design is suitable for pigs, who can exert strong force with their snout to dig or lift (29). During each familiarization session, a group was presented simultaneously with the compartments, with both doors closed. If a pig spontaneously opened a door, the door was left open for two minutes so that pigs could freely explore the inside of the compartment. At that point, any pigs still in the compartment were gently led out and a researcher closed the door in a standardized way from outside the pen, using a hook to insert the door handle back into its slot. If no pig had opened the door of a compartment for five minutes, a researcher opened that door using the hook and left the door open for two minutes, before removing any pigs inside the compartment and closing the door. This approach gave all pigs within the group multiple opportunities to open doors within and across familiarization days.

Each group then experienced the testing phase, with groups again tested consecutively in a counter-balanced order across five days. The testing phase was initially similar to the familiarization phase in that on each testing day the two compartments were simultaneously attached to the home pen of one group, and pigs could freely explore and open the doors (‘pre-testing). The goal of pre-testing was to wait until pigs lost interest in the doors and compartments, since re-attaching the compartments each day drew attention to them. Researchers monitored the pigs from a different room using a live video stream. Every ten minutes a researcher entered the room and if either of the doors were opened, they led out any pigs inside the compartments and closed the doors. Once the pigs had not touched or opened any door for ten minutes (this typically occurred within 30 min of initial exposure on each test day), a researcher entered the home pen and removed the designated trapped pig, using a pre-determined pseudo-random order, so that each pig was trapped once over the course of group testing. This began the ‘separation’ condition, during which the designated trapped pig was isolated in a separate room in a 2.8 m^2^ empty arena. The goal of the separation was to induce moderate distress in the designated trapped pig, to ensure that trapped pigs would be similarly motivated to be reunited with their group. Brief social separations can occur as part of routine husbandry on farm, but such separations are known to be stressful for pigs (28). During the separation, researchers monitored the test group remotely. If no doors were opened during the first five minutes, the designated trapped pig was carried back into the room by an experimenter and placed directly into the pre-designated test compartment from above, thus having no contact with pigs in the home pen prior to being placed in the test compartment. In the event that a pig opened any doors during the initial 5-minute separation, a researcher entered the room, closed the open door(s) and left the room. We then waited an additional five minutes before returning the designated trapped pig and placing it in the pre-designated test compartment. As a result, the total isolation time for each pig ranged from 5–11 minutes (median isolation time = 5.8 minutes).

The test trial began once the trapped pig was placed in the test compartment (‘testing’). The researcher left the room and monitored the pigs remotely using live video. During the trial, the trapped pig maintained olfactory and auditory contact with their group members at all times. In addition, the trapped pig could maintain visual and limited physical contact with group members through the mesh window located on one side of the compartment. Each test trial continued until either: i) A pig opened the door of the test compartment, releasing the trapped pig, or ii) Twenty minutes elapsed without helping, at which point a researcher entered the room and opened the test compartment door to release the trapped pig. At any point during testing, pigs could also open the door of the empty (control) compartment, but this had no influence on trial duration. If the trapped pig showed signs of distress (screaming, squeals and/or escape attempts) continuously for three minutes and was not let out by a pig, a researcher opened the door, thus ending the trial for ethical reasons. This happened only once. The pigs in the home pen were also monitored continuously via live video during each trial, and none showed evidence of distress during testing. Once the door of the test compartment was opened (either by a pig or by a researcher), the pigs were left undisturbed for an additional 15 minutes and then hormone samples were collected (see section 2.4). The compartments were then cleaned and approximately 1.5 hours after the first trial, the same group experienced a second trial, with a second pig this time trapped in the previously empty compartment. After the first group completed two trials, the second group in the cohort went through the same procedure and we alternated group testing order across test days. Within and across test days, we alternated the location of the first test compartment for each group (at the front or side of the pen), so that each group experienced both locations as test compartments with similar frequencies. At the completion of the testing phase, each pig had been trapped once, and potential helpers had between 6-9 opportunities (depending on their group size) to release different members of their social group from the test compartment.

### 2.3 Behavioural measures

Behavioural data were coded from video recordings by two trained assistants using the Observer software (version 15, Noldus, The Netherlands), with an inter-observer reliability of 91.2%. During the familiarization phase, the daily latency for each door to first be opened within each group was recorded as a measure of group task performance across familiarization days. We also recorded each pig’s daily frequency of door-opening, as a measure of task proficiency.

During the separation we recorded the identity of any pigs who opened doors, which door was opened (the designated empty or test compartment) and each pig’s frequency of door-opening. During testing, we recorded the amount of time that each potential helper (referring to all other pigs in the group besides the trapped pig), spent looking in the direction of the window of either compartment, defined as being within 10 cm of the window with their head oriented towards the window. If a pig opened the door of either compartment within 20-min, we recorded the identity of the door-opener, and their latency to open the door, based on the time at which the latch of the door was successfully lifted and the door swung open. When a potential helper released the trapped pig from the test compartment within 20 minutes, they were designated as the helper for that trial. We continued to record any door-opening of the empty compartment up to 20 minutes, to directly compare pigs’ motivation to open both compartments. We used video cameras mounted directly above the compartment to record all occurrences of squeals or screams by the trapped pig, which were identified as prolonged (> 2 sec) vocalizations with high-frequency components and amplitude modulation (30). Squeals or screams are commonly associated with negative affect in pigs (30,31). As further validation that we were recording vocalizations related to negative affect, and to test whether parts of the data analysis can be automated in future studies, we compared our manual coding of screams and squeals with analysis of vocalizations using the Stremodo software, which was developed for the automated detection of distress in pigs ((32), see also Supplementary materials). We also recorded each trapped pig’s rate of escape attempts (attempting to climb or jump against the side of the compartment) and their investigations of either the window or the door, with investigations counted whenever the snout of the trapped pig was within 5 cm of the object. For two pigs, the compartment was not entirely visible in the video recording while they were trapped, so behavioural data were analysed for the remaining n = 73 trapped pigs.

### 2.4. Hormonal measures

We assessed the physiological responses of trapped pigs by measuring changes in their concentration of sCORT, which increases in response to physical or psychosocial challenges in pigs (28). We collected saliva samples from the designated trapped pig in their home pen before they were removed from the group (pre-test), and 15 minutes after the door of the test compartment was open and the trapped pig was released (post-test). During the post-test sampling, we also attempted to collect saliva samples from the pig that helped when this could be quickly confirmed without additional video analyses. Due to the time delay between exposure to a stimulus and peak cortisol responses in pig saliva (33), the post-test samples should reflect free cortisol concentrations around the time that pigs were released. Samples were collected voluntarily from pigs using SalivaBio^®^ Infant Swabs (Salimetrics, CA, USA). Cortisol analyses were conducted at the FBN, using an ELISA kit for salivary cortisol (Demeditec Diagnostics®, GmbH, Kiel, Germany) that our lab has previously validated in pigs (28, see also supplementary materials). The inter-assay CVs of low- and high-concentration pooled controls were 9.3% and 8.3% respectively (n = 16 samples). Intra-assay CVs of low- and high-concentration pooled samples were 10.7% and 8.6% respectively.

### 2.5 Statistical analyses

Data were analysed using the R statistical software (version 4.2.1, (34), see also supplementary materials). Results are presented as medians and ranges unless otherwise specified. When test predictors in models were significant, we conducted pairwise post-hoc comparisons on marginal means, and calculated Tukey’s HSD-adjusted *p*-values (35). To measure door-opening performance during familiarization, we fit a linear mixed model (LMM) testing whether the latency for each door to first be opened was influenced by an interaction between familiarization day and location of the compartment (front or side). We included daily counter-balanced test order of each group (first or second) as a control predictor and group identity as a random effect. We also determined the proportion of pigs that successfully opened a door on each familiarization day and cumulatively across that day and previous days. To test whether pigs had a side preference to open a specific compartment during familiarization, we tested whether the proportional door-opening of each compartment was equal at the group and individual pig level using one-sample binomial tests.

To compare door-opening in helping and non-helping contexts, we fit a generalized linear mixed model (GLMM) to test for the n = 74 full trials, whether the likelihood to open either door (response) was influenced by an interaction between condition (separation or test trial) and the identity of the door (designated empty or test compartment). We included the location of each compartment (front or side), the test day (1-5), and the daily test order within a group (first or second) as control predictors, and group identity as a random effect. Then, for the subset of separations (n = 35, 47%) and test trials (n = 68, 92%) where at least one door was opened, we fit an LMM to test whether the latency to first open a door (response) was influenced by the same test and control predictors.

To test whether helping was influenced by potential helpers’ previous experience and their appraisal of the situation, we fit a GLMM with a zero-inflated negative binomial error distribution. For each potential helper (n = 7-9 pigs) in each of the 75 trials, we indicated whether that group member helped the trapped pig or not (response). We then tested how each pig’s likelihood of helping was influenced by their mean daily proficiency in door-opening during familiarization and their proportion of time spent near the window of the test compartment with their head oriented towards the window while a pig was trapped. We considered this to be a measure of social attentiveness, since from this position potential helpers should have a clear view of the trapped pig. We included control predictors for the proportion of trial time that each potential helper spent looking at the window of the empty compartment, their rate of opening either compartment during the preceding separation, and whether they opened the empty compartment during the test trial. We also included test day, daily test order within the group and the potential helpers’ relatedness to the trapped pig (sibling or non-sibling). We included group identity as a random effect.

We tested for correlations between different behavioural responses of trapped pigs using Pearsons correlation coefficients with Bonferroni corrections for multiple testing. Rates of distress vocalizations (screams or squeals) and escape attempts were correlated (see results), so we summed these two measures to obtain a combined rate of distress signals. We fit a LMM to test whether changes in sCORT concentrations in trapped pigs from pre- to post-test samples were related to their behavioural responses while trapped, for n = 70 pigs for which we had complete data on their physiological and behavioural responses to being trapped. The response was each trapped pig’s salivary cortisol concentrations. To model changes in sCORT in trapped pigs, we tested for two-way interactions between sampling context (pre- or post-trapped, representing relative changes in sCORT), and the following test predictors: i) each trapped pig’s rate of investigations to the window, where they could potentially make contact with group members, ii) their rate of distress signals (the sum of their screams, squeals and escape attempts), iii) the duration that each pig was separated from the group before being trapped and iv) the duration that each pig was trapped. As control predictors, we included each pig’s sex, the test day and daily group testing order and the time that each sample was taken. We included group and subject as random effects. We used paired t-tests and Pearson correlation coefficients to compare sCORT concentrations of trapped pigs and their helpers shortly after pigs had been released.

To test whether helpers’ responses were sensitive to the expressions of need by the trapped pig, we fit a Cox proportional hazards regression model using the Survival package (v 3.2-7; (36)). In survival analyses, latencies and likelihoods of an event of interest (in this case being released from the test compartment by a pig) are combined into a hazard ratio (HR), with an HR above one indicating a covariate that is associated with a greater likelihood and shorter latency for the event to occur. Trials in which pigs were released by humans were right-censored, to indicate that the event of interest did not occur during that pig’s observation period. To compare behavioural data that varied over time while ensuring that we did not violate the proportional hazards assumptions, we stratified each trapped pig’s behavioural data at one-minute intervals and removed any intervals of less than 60-sec at the beginning of each pig’s trapped period. We also excluded from analysis each pig’s behaviour during the last 10-secs before they were released, to avoid committing “immortal biases” (37) by using information from the future to predict the outcome. After removing n = 15 trapped pigs who were released in under one minute, and the n = 2 trapped pigs mentioned above whose behavioural data could not be analysed due to limited test compartment visibility, we ran the survival analysis on n = 58 trapped pigs. We included each trapped pig’s number of investigations to the mesh window per minute, which could reflect motivation to acquire information about the social group or to make social contact. The number of distress signals (screams, squeals or escape attempts) per minute could not be transformed to approximate normality. Therefore, we included the presence or absence of distress signals at one-minute intervals as a binomial predictor. We included as control predictors group identity, sex, the counter-balanced location of the test compartment (at the front or the side of the pen), the group testing order and the time when the trapped pig was first placed in the compartment. Diagnostic tools indicated that the proportional hazards assumptions were not violated (global hazard ratio, χ2 = 22.07, df = 21, p = 0.40). The Concordance Index (CI) was 0.69, indicating good predictive discrimination of the covariates. All data and code related to this study are available on-line at: https://doi.org/10.5281/zenodo.8096753.

## 3. Results

### 3.1 Door-opening during familiarization

Despite having no exposure or training in the task beforehand a median of 81 (range = 77–93)% of pigs in each group opened doors on each familiarization day and by the end of familiarization all but one of the 75 pigs had opened a door at least once (see **Fig. S3**). The interaction between location of compartment and familiarization day tended to influence door-opening latency (LMM, *F*_4,72_ = 2.50, p = 0.051, see **Fig. S4 and Table S1**). On day 3, pigs were faster to open the side compartment than the front compartment (0.10 (0.01–0.48) min vs 1.35 (0.04–3.48) min, p = 0.004). Door-opening latencies on other days were not affected by the location of the compartment (p values > 0.19). Groups were also faster to open the side compartment on day 3 compared to day 1 (0.70 (0.35–7.26) min) and day 2 (0.75 (0.18–1.45) min, p-values < 0.027, see **Fig. S4**). There were no differences in latencies to open the front compartment across familiarization days (p-values > 0.17). Pigs varied in their door-opening proficiency with 2 (0–11) daily door openings per pig, but only four (5%) showed evidence for a location preference for door-opening, with three opening the side door more frequently (p-values < 0.02), and one opening the front door more frequently (p = 0.035, see **Table S2**). At the group level, there was no preference to open one of the two doors more frequently during familiarization (binomial tests, p-values > 0.18, see **Table S2**).

### 3.2 Door-opening in non-helping and helping contexts

During the separation and testing conditions, potential helpers had opportunities to perform the same door-opening behaviour in non-helping contexts, by opening either compartment during the separation, before the trapped pig was placed in the designated test compartment, or by opening the empty compartment during the test trial. During the separation, 37 pigs (49% of potential helpers) opened doors, during 36 trials (48% of trials), with a median of 0 (range = 0-5) door-openings per separation. This accounted for the variable duration of isolation time (from 5 to 11 min) required to meet the criteria that no door had been opened for five minutes before a test trial began. During the test trials, 85% (n = 64) of the trapped pigs were helped by a group member, with a latency of 2.2 (0.2–17) min to be released (See **Fig. S5**). Of the additional 11 pigs, ten were released by a researcher after the 20-minute trial duration ended and one trial was aborted and the pig released for ethical reasons after exhibiting distress signals for three minutes without being helped. In total, n = 33 pigs (44% of potential helpers) helped a trapped pig, by being the quickest to first open the door of the test compartment, and each helper helped a median of two trapped pigs (range = 1–4 pigs, or 11–57% of their group members). During 68% (n = 50) of the full n = 74 test trials, n = 28 pigs (37% of potential helpers), opened the door of the empty compartment within the 20-min maximum trial duration, with a latency of 3.9 (0.8–18.8) min to open the empty compartment door.

The likelihood of door opening was influenced by an interaction between the condition and the compartment (est ± SE = 1.50 ± 0.58, p = 0.008, see **Table S3** and **Fig. S6**). Pigs opened the test compartment during 85 (63–100)% of test trials within their group, while opening the empty compartment during 68 (40–100)% of test trials, p = 0.006). In contrast, there were no differences in the likelihood of opening the two compartments during the separation (21 (0–43)% vs. 30 (10–57)% of separations, p = 0.450). The latency to first open a door was also influenced by an interaction between condition and compartment (est ± SE = -0.97 ± 0.36, p = 0.012, see **Table 1** and **Fig. 1**). Pigs were quicker to open the test compartment than the empty compartment during test trials (p< 0.001), but did not differ in their latencies to open either of the compartments during the preceding separation period (p = 0.699, **see Fig. 1**). There was no effect of test day, test order within a group, or location of each compartment (at the front or side of the pen), on the likelihood or latency to open compartments (see **Table 1** and **Table S3**).

**Figure 1.**
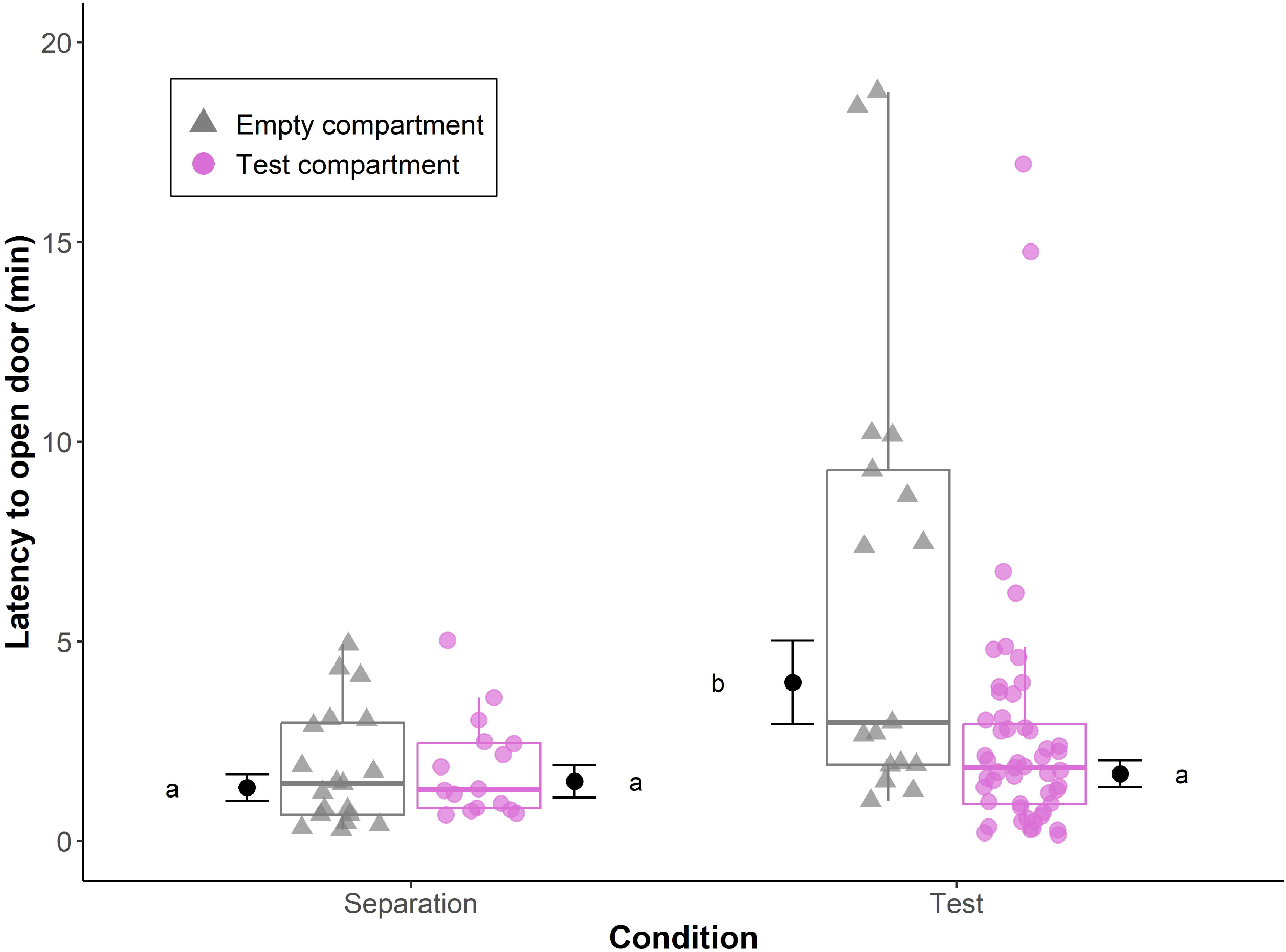
Latencies to open compartments during test trials, when a pig was trapped in the test compartment, and during the separation, when both compartments were empty. Data points are from separations and trials when at least one door was opened. Box plots indicate medians and 25-75% interquartile ranges. Estimated marginal means and SEs are also indicated, with results of pair-wise post-hoc comparisons (Tukey’s HSD) visualized using a compact letter display. Different letters indicate significant differences. For full model results see Table 1.

**Table 1.**
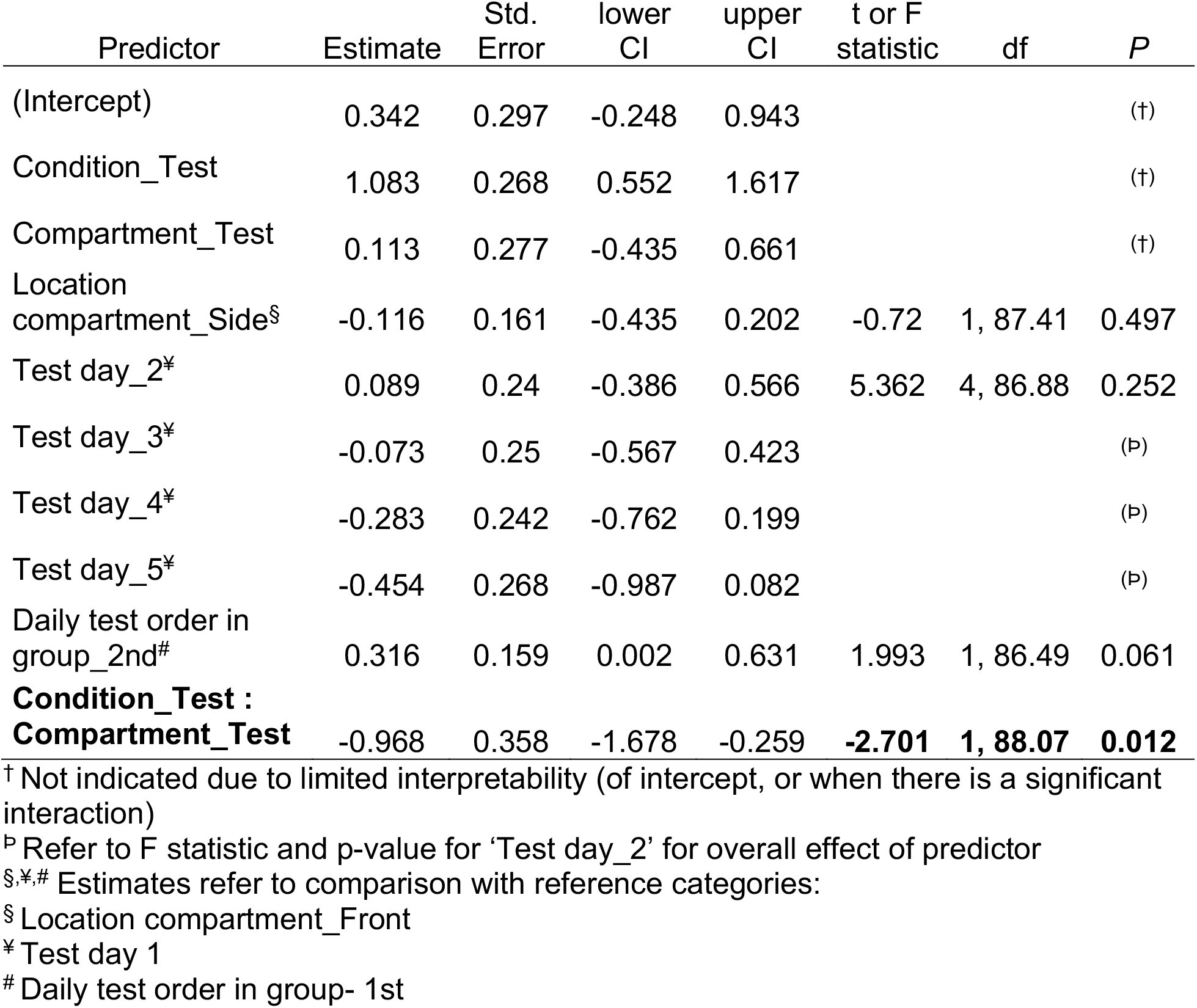
Results of a LMM testing for an influence of condition (separation vs test trial) and identity of compartment (designated empty or test compartment) on the latency for pigs to first open a door. Significant predictors are indicated in bold.

### 3.3. Behavioural and physiological responses of helpers and influences on helping

While pigs were trapped, potential helpers spent longer looking in the direction of the test compartment window (7.7 (0.1–59.6) sec per trial min) than they did looking in the direction of the empty compartment window (0.2 (0–4.4) sec per trial min, Wilcoxon signed-rank test, n = 75, Z = 8.89, p < 0.001). Potential helpers who spent a greater proportion of time looking in the window of the test compartment while a pig was trapped inside were more likely to help the trapped pig (GLMM, est ± SE = 0.51 ± 0.08, p < 0.001, see **Table 2** and **Fig. 2**). There was no relationship between proportion of time spent looking in the window of the empty compartment and helping (see Table 2). Pigs who were more proficient in door-opening across familiarization days were also more likely to help their group members (est ± SE= 0.34 ± 0.13, p= 0.008). However, a potential helper’s rate of door opening during the preceding separation, and their likelihood of opening the empty compartment during the test period, did not predict helping (see Table 2). The potential helper’s relatedness to the trapped pig, test day and daily test order within a group also did not influence the likelihood of helping a trapped group member.

**Figure 2.**
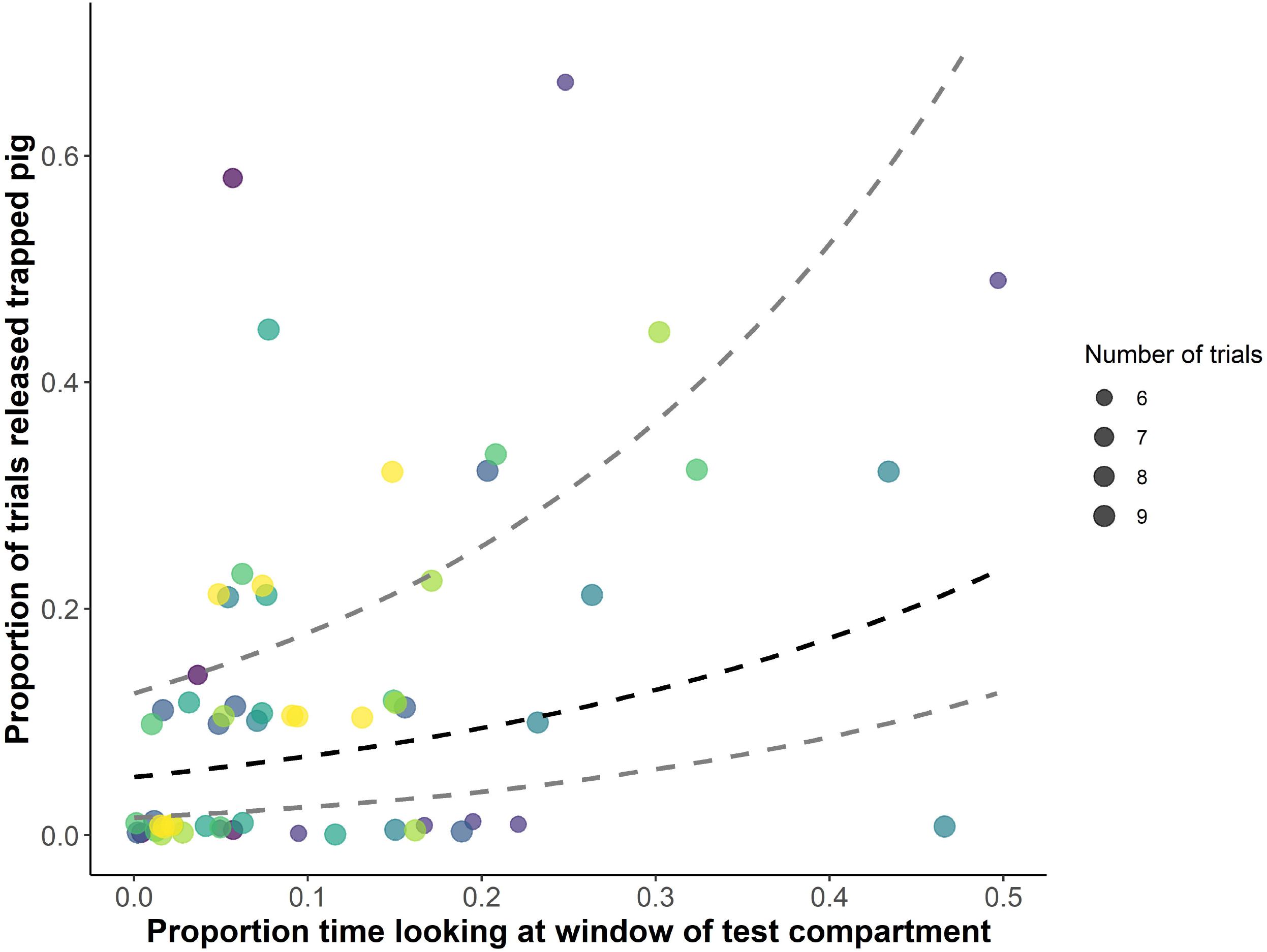
Relationship between potential helpers’ proportional time spent looking at the window of the test compartment while a pig was trapped and their likelihood of helping, averaged across their 6–9 exposures to trapped group members. Colours indicate different social groups. The dashed lines depict the estimate and 95% CI from the fitted model, with control predictors centred around 0. For full model results see Table

**Table 2.**
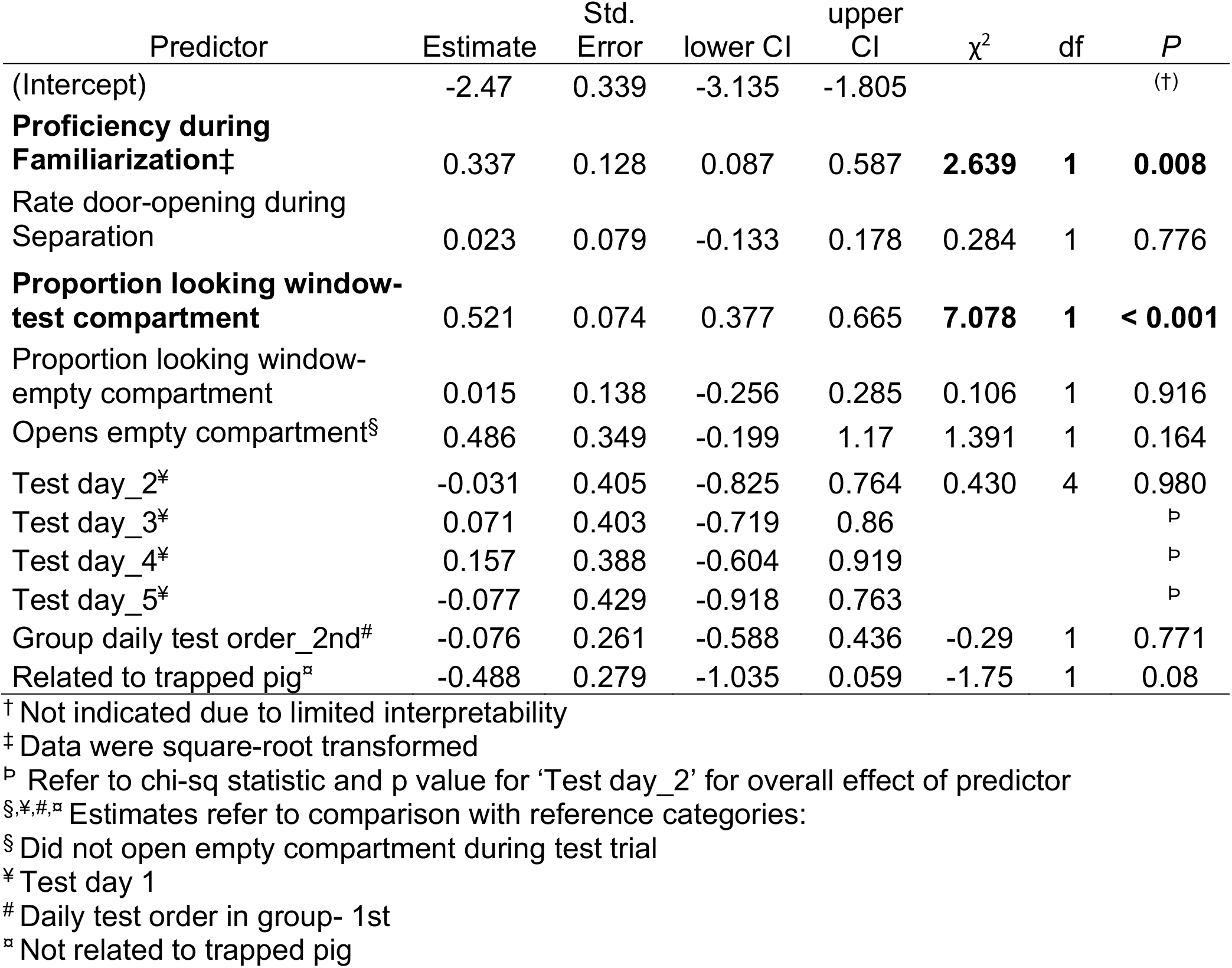
Results of a zero-inflated negative binomial GLMM predicting influences on helping responses. Significant predictors are indicated in bold.

In 67% of the helping trials (n = 43 out of 64), the helper initiated nose-to-nose or nose-to-body contact with the previously trapped pig within one minute after opening the test compartment door. In 81% of helping trials (n = 52), the helper also entered the test compartment within one minute of helping. In 54% (n = 28) of these trials, the previously trapped pig had already left the compartment, so entering the compartment did not lead to social contact with the trapped pig.

For n = 50 trials where we were able to quickly identify the helper without requiring additional video analyses, we obtained saliva samples from the helper as well as the released pig fifteen minutes after helping. Helpers had lower sCORT concentrations than their released group mate (Mean ± SD = 3.9 (± 1.9) ng/mL vs 7.3 (± 2.4) ng/mL, *t*(49) = 8.14, p < 0.001, see **Fig. S7**), and there was no correlation between sCORT concentrations of released pigs and their helpers shortly after release (Pearson, r(48)= = 0.07, p = 0.63).

### 3.4 Behavioural and physiological responses of trapped pigs and influences on being helped

Of the 73 trapped pigs for whom behavioural data could be analysed from video recordings, 72 investigated the window and 71 investigated the door. Trapped pigs investigated the window 4.3 (0 - 19) times per minute trapped, and they investigated the door 1.6 (0-11.6) times per minute. Rates of investigations to the window and door were positively correlated (r(71) = 0.80, p < 0.001). The majority of trapped pigs (74%, n = 54) attempted to escape and 44% of trapped pigs (n = 32) emitted screams or squeals while trapped. Trapped pigs’ rates of distress vocalizations and escape attempts were positively correlated with each other (r(71) = 0.34, p = 0.02), but not with either of the trapped pigs’ investigatory behaviours (r(71) < 0.22, p values > 0.36). We therefore summed rates of distress vocalizations and escape attempts to create a combined measure of distress signals.

For n = 72 pigs with sufficient saliva volumes to measure sCORT before and after being trapped, 87.5% (n = 63) exhibited increases in sCORT in their post-trapped samples (Mean ± SD = 7.3 (± 2.7) ng/mL), compared to their samples before being trapped (Mean ± SD = 4.7 (± 2.0) ng/mL, t(71) = 7.97, p < 0.001). Pigs with greater rates of distress signals while trapped tended to have greater increases in sCORT (interaction: est ± SE = -0.124 ± 0.061, p = 0.045, **Table S4** and **Fig. S8**). Changes in sCORT concentrations while trapped were not related to a pig’s duration of separation, their duration of time trapped or other control predictors (see Table S4).

The Cox proportional hazards model indicated that trapped pigs who exhibited more distress signals were 2.4 times more likely and quicker to be subsequently released, compared with pigs who exhibited fewer distress signals (χ^2^ = 4.65, df = 1, p = 0.031, **Table 3**, **Fig. 3 and Supplementary videos**). The rate of window investigations by trapped pigs did not influence helping outcomes (see Table 3). There were also significant differences among groups in the likelihood that trapped pigs would be helped (χ^2^ = 16.26, df = 7, p = 0.023, see Table 3). The other control predictors did not influence the probability of being helped.

**Figure 3.**
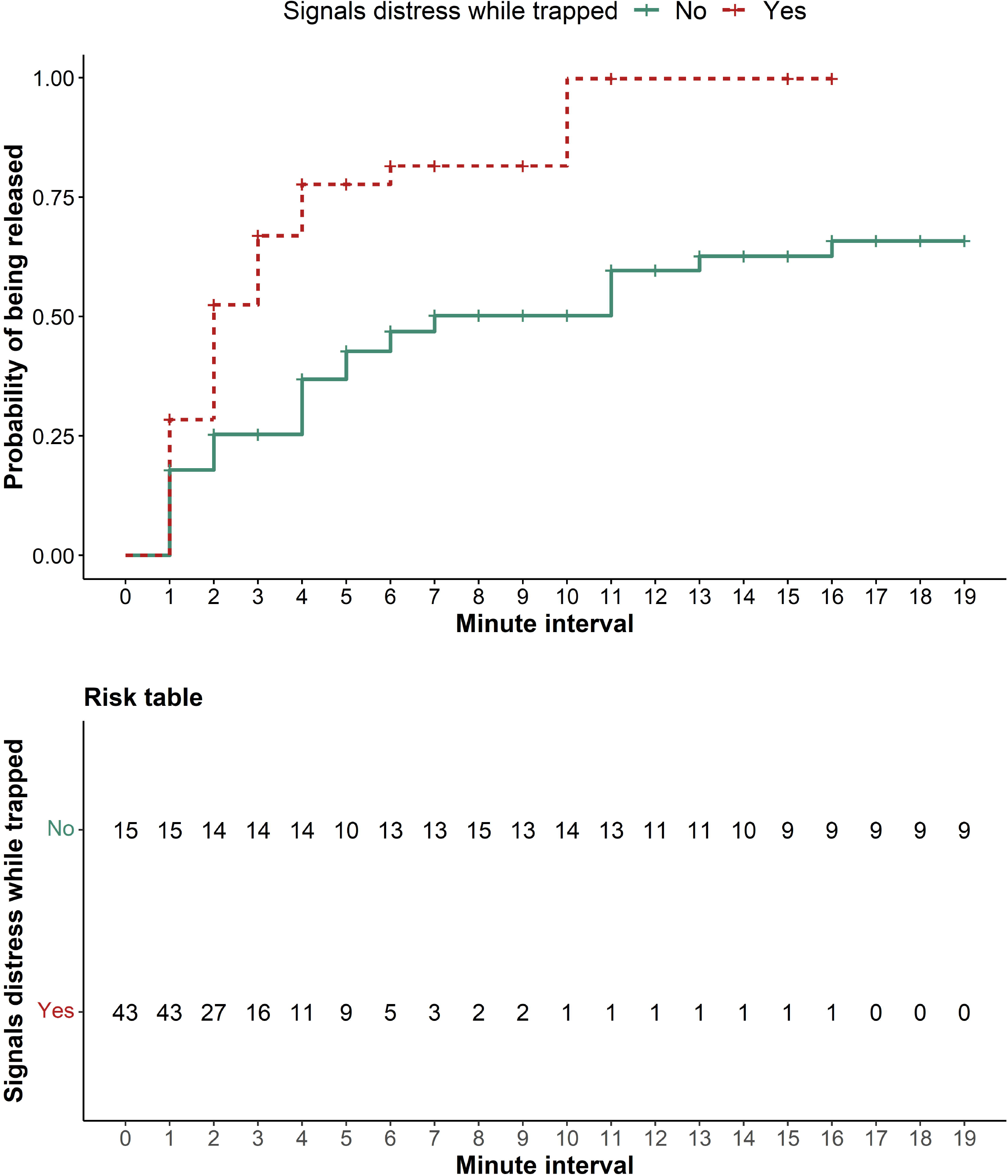
Survival curves showing that pigs who gave distress signals at any given minute interval while trapped were more likely to be released than pigs who did not (Hazard ratio = 2.4, p = 0.03). For full results, see Table 3.

**Table 3.** Results of a survival analysis predicting the probability for trapped pigs to be helped. Hazard ratios are indicated by the exponentiated coefficients. Significant predictors are indicated in bold.

| Predictor | coef | exp(coef) | se(coef) | robust<br>se | $\chi^2$ | df | *P* |
| --- | --- | --- | --- | --- | --- | --- | --- |
| **Signals distress_Yes<sup>†</sup>** | 0.86 | 2.37 | 0.41 | 0.40 | **4.65** | **1** | **0.031** |
| Rate window investigations<sup>‡</sup> | -0.18 | 0.84 | 0.22 | 0.20 | 0.81 | 1 | 0.368 |
| Trapped pig sex_Female<sup>§</sup> | 0.05 | 0.95 | 0.34 | 0.33 | 0.03 | 1 | 0.868 |
| Time in compartment | 0.03 | 1.03 | 0.16 | 0.14 | 0.04 | 1 | 0.836 |
| Location<br>compartment_Side<sup>¥</sup> | 0.39 | 1.48 | 0.34 | 0.34 | 1.37 | 1 | 0.242 |
| Test order in group | 0.15 | 1.16 | 0.16 | 0.16 | 0.91 | 1 | 0.341 |
| **Group identity_1B<sup>#</sup>** | 1.46 | 4.31 | 0.92 | 0.80 | **16.26** | **7** | **0.023** |
| **Group identity_2A<sup>#</sup>** | -0.14 | 0.87 | 0.67 | 0.72 |  |  | (<sup>Ⓟ</sup>) |
| **Group identity_2B<sup>#</sup>** | 1.00 | 2.72 | 0.62 | 0.65 |  |  | (<sup>Ⓟ</sup>) |
| **Group identity_3A<sup>#</sup>** | 0.51 | 1.66 | 0.64 | 0.69 |  |  | (<sup>Ⓟ</sup>) |
| **Group identity_3B<sup>#</sup>** | 0.73 | 2.07 | 0.61 | 0.62 |  |  | (<sup>Ⓟ</sup>) |
| **Group identity_4A<sup>#</sup>** | -0.04 | 0.97 | 0.62 | 0.67 |  |  | (<sup>Ⓟ</sup>) |
| **Group identity_4B<sup>#</sup>** | 1.33 | 3.78 | 0.65 | 0.60 |  |  | (<sup>Ⓟ</sup>) |

<sup>†,§,¥,#</sup> Estimates refer to comparison with reference categories:
<sup>†</sup> Trapped pig did not signal distress during the preceding interval
<sup>§</sup> Trapped pig sex\_Male
<sup>¥</sup> Location compartment\_Front
<sup>#</sup> Group Identity\_1A
<sup>‡</sup> Data were square-root transformed
<sup>Ⓟ</sup> Refer to chi-square statistic and p value under “Group Identity\_1B” for the over-all effect of the predictor

## 4. Discussion

Pigs spontaneously opened doors without training, and subsequently preferred to open doors to free a trapped group member than to open doors to empty compartments. Pigs who directed more attention towards the window of the compartment where a pig was trapped were quicker to help, and distress signals from trapped pigs increased their chances and decreased their latency to be helped. Our results support anecdotal evidence for spontaneous rescue behaviour in wild boars (*Sus scrofa*), the ancestor of domestic pigs (7). We avoid using the term ‘rescue’ here, since not all trapped pigs exhibited signs of distress, and helping did not necessarily require a great cost or risk, which are two of the criteria suggested by Hollis and Nowbahari (38) to define rescue behaviour.

There was considerable variation in the likelihood that pigs helped trapped group members, and in the latency for trapped pigs to be released, which enabled us to investigate the factors mediating helping behaviour. Roughly half (44%) of the pigs in each group helped others during the test phase. However, given that only one pig could be a helper per trial, our group-testing paradigm may also underestimate the propensity for helping behaviour at the population level. Pigs who opened doors at greater rates during familiarization were more likely to help others, suggesting that their ability or willingness to solve this novel task played a role in their helping behaviour.

Throughout the testing phase, pigs also opened doors in non-helping contexts, by opening either door during the separation (49% of potential helpers), or by opening the empty door during a test trial (37% of potential helpers). However, few of these individuals (24% or 9 of 37 pigs who opened doors during the separation phase and 32% or 9 of 28 pigs who opened the empty door during a test trial) also opened the door of the test compartment to help the trapped pig. As a result, pigs’ door-opening in non-helping contexts during the separation or test trial did not influence their likelihood of opening the test compartment to help the trapped pig. Across the majority of familiarization days and during separations, pigs had no preference to open one of the two compartments. In contrast, during test trials pigs were more likely and quicker to open the compartment containing a trapped group member than the empty compartment, suggesting that door-opening during test trials was goal-oriented.

While a pig was trapped, potential helpers spent more time looking at the window of the test compartment than they spent looking at the window of the empty compartment, and the proportion of time spent looking at the test compartment window was the strongest predictor of helping. Pigs who spent more time oriented towards the test compartment window at close range may have been visually assessing the situation of the trapped pig, which would be consistent with an other-oriented response (26). However, the majority of trapped pigs produced vocal and locomotor distress signals that should have been readily detectable without looking through the window, and the helpers’ behaviour is also consistent with more selfish explanations. As an alternative explanation, it is likely that the movements and vocalizations of the trapped pig acted as salient features to attract attention to the test compartment and window, and distressed pigs likely produced more salient signals that were more effective in attracting attention. Potential helpers may have then looked in the window to gather information about the situation for themselves (e.g. 13,16). As a result, potential helpers would also be closer to the door and handle, increasing the chances of door-opening through an effect of local enhancement. But if local enhancement explained helping behaviour, we would expect the majority of potential helpers to be similarly interested in and attentive to highly salient cues from trapped pigs in distress, which was not the case (see Fig. 2 and supplementary videos).

Distress signals given by the trapped pig, in the form of screams, squeals and escape attempts, increased the pig’s subsequent chances of being released by a 2.4 odds ratio. This evidence is consistent with the interpretation of helping behaviour as a flexible response that is appropriate to the situation of the trapped pig, which is another criteria for targeted helping (26,39). It is important to note that distress signals were neither necessary nor sufficient for helping to occur. Of the n = 15 pigs who were released in under one minute and thus could not be included in the survival analysis, 67% (n = 10) did not produce measurable distress signals before being released. That helpers sometimes quickly freed these pigs counters arguments that spontaneous helping may merely be motivated by selfish needs to alleviate the source of aversive distress signals (24). However, it also raises the question of whether all trapped pigs were in need of help (16). In addition, of the eleven pigs who were released by humans, nine (82%) gave distress signals at some point while trapped, although (with the exception of the aborted trial), these pigs gave distress signals at lower rates compared with pigs who were released by conspecifics, as indicated by the survival analysis.

A primary concern of animal helping paradigms is that appropriate controls are often not provided to test whether helping behaviour is motivated by self-interest. One of the main examples is when helping provides potentially rewarding social interactions for helpers. Indeed there is ample evidence that helping decreases when post-helping social interactions are prevented (14,18,22), although such designs may also make it more difficult for potential helpers to understand the causal effects of their actions. One of the main advantages of testing animals in stable social groups in their home environment is that this reduces the likelihood that helping is due to the desire for social reinforcement more generally. Moreover, pigs were equally likely to help their siblings whom they had known since birth as they were to help unrelated group members whom they had known for a shorter time, which argues against helping motivated by social preferences. We cannot rule out the possibility that the helper may have had an interest in interacting with the specific trapped individual, which is supported by the fact that the majority of helpers initiated social contact with the previously trapped pig within a minute of helping. More work is thus needed to determine whether dyadic social preferences or dominance relationships influence helping behaviour.

In the majority of cases (81%), pigs entered the test compartment immediately after helping, although in most of these cases the trapped pig had already left the compartment. In some rodent helping paradigms, similar proportions of helpers entered a tube where conspecifics had previously been restrained (19,40). In those cases, it was hypothesized that rodents may be motivated to enter the restraint tube because it is novel (40), and may even prefer it to a barren testing arena (41). Such explanations are unlikely in our paradigm, since pigs were tested in their home environment, had been familiarized to the compartments prior to testing, and had equal opportunities to access the empty compartment, but were less likely and slower to do so. It is more likely that helpers entered the test compartment to gather relevant information for themselves after witnessing the trapped pig, which is consistent with selfish motivations.

Wild boars reside in stable, cohesive social groups and even brief separations from their social group can induce behavioural and physiological signs of stress in domestic pigs (28,33). Either the separation alone, or the separation in combination with being in the test compartment are likely to have been stressful for pigs. The majority of pigs had increases in sCORT concentrations while trapped, and trapped pigs with greater increases in sCORT exhibited more behavioural responses consistent with distress. However, helpers did not show corresponding increases in sCORT concentrations, as expected if they experienced emotional contagion of distress. In contrast, their sCORT around the time of helping was similar to baseline cortisol concentrations of pigs prior to being trapped. Although emotional contagion has been shown in pigs in non-helping contexts (25), it does not appear to explain helping behaviour in this case. In fact, in rodent studies, potential helpers who showed the greatest stress-reactivity to trapped conspecifics were less likely to help than those who showed more moderate arousal (21). Indeed, a moderate level of emotional arousal, consistent with emotional control regulation, is one of the criteria for identifying empathic targeted helping in animals (26). The cortisol responses of helpers in this study were more consistent with low rather than moderate arousal, suggesting that helpers were not stressed by witnessing trapped pigs. This is consistent with a previous study in which juvenile pigs did not show emotional contagion of fear after hearing playbacks of distress vocalizations from unfamiliar pigs (42). Further research should compare the physiological responses of helpers and non-helpers to better assess how arousal relates to helping.

Given the promising evidence that potential helpers who attended more closely to the test compartment window were more likely to help, follow-up research should test whether a broader range of communicative signals by trapped pigs may mediate helping. Two likely candidates are low-frequency grunt vocalizations and nose-to-nose contacts, both of which are common socio-communicative behaviours in pigs (43). Grunts and nose-to-nose contacts were observed during helping trials but could not be systematically investigated with our recording equipment. Helping paradigms in which individuals in need can express a broader range of behavioural responses are important for clarifying the role of social communication in influencing helping and for further distinguishing between self- and other-oriented explanations for helping behaviour in animals.

## 5. Conclusion

This group-testing paradigm offers a novel experimental approach to systematically study the proximate factors mediating spontaneous helping in a broad range of social species, including farm animals. Applying the framework suggested by Pérez-Manrique & Gomila (26) for evaluating comparative evidence for empathy, our results provide partial support for targeted helping in that: i) helpers exhibited visual assessment of the situation, ii) trapped pigs who signalled more distress were more likely and quicker to be helped and iii) helping improved the situation for trapped pigs by reuniting them with their social group. However, the lack of evidence for physiological arousal in helpers argues against an other-oriented response (26). We also point out alternative explanations based on local enhancement and selfish motivations to enter the test compartment that can more readily explain why the majority of helpers entered the compartment after helping, and why a small percentage of trapped pigs were helped without showing obvious signs of distress. Thus, follow-up studies are needed to clarify the motivations underlying spontaneous helping in pigs.

## Acknowledgements

We thank V. Hörsch, B. Sobczak and E. Normann for their assistance with running the experiments and collecting saliva samples. We thank P. Müntzel for conducting the laboratory analyses. We are grateful to V. Hörsch and M. Wagenknecht for their help with behavioural coding of videos. We thank M. Zenk for her help with the coordination of this project and D. Ameling, H. Asmus, F. Hintze and H. Schumann for their daily care of the pigs at the Experimental Pig Facility of the FBN. We are especially grateful to T. Therneua, for his repeated advice that helped us to improve our survival analysis, and for helpful feedback from two anonymous reviewers that inspired us to conduct several additional analyses.

