## Supplementary Materials for "Spontaneous helping in pigs is mediated by helper’s social attention and distress signals of individuals in need"

**Supplementary materials for the manuscript:** Moscovice, L. R., Eggert, A., Manteuffel, C. and Rault, J- L. (2023): *Spontaneous helping in pigs is mediated by helper's social attention and distress signals of individuals in need. Proc. Royal Soc. B. DOI:* 10.1098/rspb. 2023.0665

#### Supplementary Text

#### Methods

##### 2.1 Animals and husbandry

After birth, piglets were housed with their mothers in 6 $m^2$ farrowing pens. Piglets had *ad libitum* access to water and were offered dry food (HAKRA-Immuno-G; Una Hakra, Hamburg, Germany) beginning from 14 days of age in addition to milk from their mothers. Piglets received tattoos for identification at 1 day of age and ear tags with unique numbers at 27 days of age. Piglets were not subjected to tail docking, teeth clipping, castration or other invasive procedures, and none of the study subjects were cross-fostered to other litters, which can occur in farm practice. Throughout testing, indoor lighting in the room was controlled on a timer, with lights on from 7:00-15:00 hr. Pigs also received natural light through a window. Food (Porcistart, Trede und von Pein GmbH, Damfleth, Germany) and water were provided *ad libitum*, and straw and an enrichment object (Easyfix® Luna 68, Ballinasloe, Ireland) were provided daily. Room temperature was maintained between 20-22°C.

##### 2.3 Behavioural Measures

As further validation that we were recording vocalizations related to negative affect, and to test whether parts of the data analysis can be automated in future studies, we compared our manual coding of screams and squeals with analysis of vocalizations using the Stremodo software, which was developed for the automated detection of distress in pigs (Schön et al., 2004). Stremodo detects distress signals in 100 ms time windows of audio recordings without considering the context. To facilitate comparison with human classifications of vocalizations, a post-processing routine automatically detected the beginning and end of a vocalisation based on the loudness of the audio data. Then this vocalisation as a whole was marked as a stress event regardless of its duration, if Stremodo identified one or more of its segments as stressful. Measures of distress vocalizations based on manual coding of screams or squeals and the automated Stremodo analysis were positively correlated ($r(71) = 0.54$, $p < 0.001$). The lack of a stronger correlation is likely related to interference from additional noises in the room, which the Stremodo algorithm may have sometimes falsely classified as screams or squeals. We used data from manually-coded distress vocalizations in subsequent analyses.

**2.4 Hormonal measures**

Sample collection took one to two minutes per pig. Swabs were placed in polypropylene tubes on ice immediately after collection and were centrifuged within 30 minutes of collection at 4° C (15 min, 2,000 × g). The eluent was stored at -80°C until hormone analyses.

**2.5 Statistical Analyses**

As a general framework for the statistical models, for continuous response variables, we fit linear mixed models (LMMs) and generalized linear mixed models (GLMMs) using the R package `lme4` (Bates et al., 2015). For zero-inflated negative binomial responses, we fit a GLMM using the R package `glmmTMB` (Brooks et al., 2017). For LMMs, we tested for significance levels of the fixed effect predictors using t-tests with the Satterthwaite approximation (Luke, 2017) in the function lmer of the package lmerTest (v 3.1-3, Kuznetsova et al., 2017) and a model fitted with restricted maximum likelihood. For all models, continuous response and predictor variables were square-root or log-transformed when necessary to approximate normal distributions. Continuous predictors were z-transformed to facilitate interpretability of the results. We used likelihood ratio tests to compare the full model to a reduced model excluding the test predictors, and present results when the model differed significantly from the reduced model. We assessed model performance by checking for normality of residuals (for LMMs), over-dispersion (for GLMMs) and collinearity of predictors (for all models) using the `performance` package (Lüdecke et al., 2021). We used likelihood ratio tests to compare the full model to a reduced model excluding the test predictors, and present results when the model differed significantly from the reduced model.

**Supplemental Table S1.** Results of a LMM testing for an influence of familiarization day (1-5) and location of compartment (front or side of pen) on the latency for pigs to first open a door. Predictors that tend towards significance are indicated in bold.

| Predictor | Estimate | Std. Error | lower CI | upper CI | t or F statistic | df | *P* |
|---|---|---|---|---|---|---|---|
| (Intercept) | 4.509 | 0.484 | | | | | (†) |
| Location Compartment_Side | -0.353 | 0.643 | -1.630 | 0.923 | | | (†) |
| Familiarization day_2 | -1.558 | 0.643 | -2.834 | -0.282 | | | (†) |
| Familiarization day_3 | -0.659 | 0.643 | -1.936 | 0.617 | | | (†) |
| Familiarization day_4 | -1.166 | 0.643 | -2.443 | 0.110 | | | (†) |
| Familiarization day_5 | -1.319 | 0.643 | -2.595 | -0.042 | | | (†) |
| Group daily test order | -0.318 | 0.289 | -0.894 | 0.265 | -1.019 | 75.76 | 0.311 |
| **Location Compartment_Side: Familiarization day_2** | 1.270 | 0.909 | -0.535 | 3.075 | **2.500** | **4, 71.86** | **0.051** |
| **Location Compartment_Side: Familiarization day_3** | -1.716 | 0.909 | -3.521 | 0.090 | | 71.86 | (ᵖ) |
| **Location Compartment_Side: Familiarization day_4** | -0.447 | 0.909 | -2.252 | 1.358 | | 71.86 | (ᵖ) |
| **Location Compartment_Side: Familiarization day_5** | 0.328 | 0.909 | -1.477 | 2.133 | | 71.86 | (ᵖ) |

† Not indicated due to limited interpretability (of intercept, or when there is a significant interaction)
ᵖ Refer to F statistic and p-value for ‘Location Compartment_Side: Familiarization day 2’ for overall effect of interaction

**Supplemental Table S2.** Results of binomial tests for preferences to open front and side compartments at the group (a) and individual (b) level during familiarization days. Four pigs who exhibited side preferences are indicated in bold.

**a)**

| Group Identity | Opening-Front compartment | Opening-Side compartment | binom.p |
|---|---|---|---|
| Group1_A | 40 | 52 | 0.25 |
| Group1_B | 78 | 62 | 0.20 |
| Group2_A | 75 | 78 | 0.87 |
| Group2_B | 73 | 91 | 0.18 |
| Group3_A | 85 | 69 | 0.23 |
| Group3_B | 94 | 89 | 0.77 |
| Group4_A | 67 | 66 | 1.00 |
| Group4_B | 88 | 84 | 0.82 |

**b)**

| Pig Identity | Front | Side | binom.p |
|---|---|---|---|
| Group1_A_2 | 24 | 23 | 1.00 |
| Group1_A_3 | 2 | 5 | 0.45 |
| Group1_A_4 | 6 | 5 | 1.00 |
| Group1_A_5 | 3 | 1 | 0.63 |
| Group1_A_6 | 3 | 1 | 0.63 |
| **Group1_A_7** | **2** | **16** | **0.00** |
| Group1_A_8 | 0 | 1 | 1.00 |
| Group1_B_1 | 6 | 2 | 0.29 |
| Group1_B_2 | 9 | 12 | 0.66 |
| Group1_B_3 | 12 | 9 | 0.66 |

| | | | |
|---|---|---|---|
| Group1_B_4 | 15 | 11 | 0.56 |
| Group1_B_5 | 15 | 11 | 0.56 |
| Group1_B_8 | 8 | 6 | 0.79 |
| Group1_B_9 | 12 | 8 | 0.50 |
| Group2_A_1 | 8 | 4 | 0.39 |
| Group2_A_10 | 4 | 9 | 0.27 |
| Group2_A_2 | 7 | 11 | 0.48 |
| Group2_A_3 | 8 | 6 | 0.79 |
| Group2_A_4 | 15 | 14 | 1.00 |
| Group2_A_5 | 20 | 23 | 0.76 |
| Group2_A_6 | 3 | 5 | 0.73 |
| Group2_A_7 | 3 | 3 | 1.00 |
| Group2_A_8 | 1 | 0 | 1.00 |
| Group2_A_9 | 6 | 3 | 0.51 |
| Group2_B_1 | 7 | 2 | 0.18 |
| Group2_B_10 | 16 | 21 | 0.51 |
| Group2_B_2 | 7 | 7 | 1.00 |
| Group2_B_3 | 5 | 5 | 1.00 |
| Group2_B_4 | 3 | 0 | 0.25 |
| Group2_B_5 | 6 | 14 | 0.12 |
| Group2_B_6 | 5 | 6 | 1.00 |
| **Group2_B_7** | **7** | **20** | **0.02** |
| Group2_B_8 | 5 | 5 | 1.00 |
| Group2_B_9 | 12 | 11 | 1.00 |
| Group3_A_1 | 16 | 10 | 0.33 |
| Group3_A_10 | 4 | 4 | 1.00 |
| Group3_A_2 | 12 | 11 | 1.00 |
| Group3_A_3 | 9 | 3 | 0.15 |
| Group3_A_4 | 6 | 5 | 1.00 |
| Group3_A_5 | 9 | 7 | 0.80 |
| Group3_A_6 | 4 | 6 | 0.75 |
| Group3_A_7 | 6 | 9 | 0.61 |

| | | | |
|---|---|---|---|
| Group3_A_8 | 15 | 10 | 0.42 |
| Group3_A_9 | 4 | 4 | 1.00 |
| Group3_B_1 | 7 | 6 | 1.00 |
| Group3_B_10 | 4 | 2 | 0.69 |
| Group3_B_2 | 16 | 6 | 0.05 |
| Group3_B_3 | 2 | 0 | 0.50 |
| Group3_B_4 | 20 | 12 | 0.22 |
| Group3_B_5 | 12 | 8 | 0.50 |
| Group3_B_6 | 13 | 22 | 0.18 |
| Group3_B_7 | 3 | 10 | 0.09 |
| **Group3_B_8** | **2** | **12** | **0.01** |
| Group3_B_9 | 15 | 11 | 0.56 |
| Group4_A_1 | 13 | 14 | 1.00 |
| Group4_A_10 | 3 | 5 | 0.73 |
| Group4_A_2 | 9 | 6 | 0.61 |
| Group4_A_3 | 9 | 13 | 0.52 |
| Group4_A_4 | 10 | 3 | 0.09 |
| Group4_A_5 | 3 | 2 | 1.00 |
| Group4_A_6 | 8 | 5 | 0.58 |
| Group4_A_7 | 6 | 6 | 1.00 |
| Group4_A_8 | 0 | 4 | 0.13 |
| Group4_A_9 | 6 | 8 | 0.79 |
| Group4_B_1 | 9 | 11 | 0.82 |
| Group4_B_10 | 8 | 4 | 0.39 |
| Group4_B_2 | 8 | 6 | 0.79 |
| Group4_B_3 | 19 | 22 | 0.76 |
| Group4_B_4 | 6 | 6 | 1.00 |
| Group4_B_5 | 11 | 13 | 0.84 |
| Group4_B_6 | 9 | 5 | 0.42 |
| Group4_B_7 | 5 | 10 | 0.30 |
| Group4_B_8 | 0 | 4 | 0.13 |
| **Group4_B_9** | **12** | **3** | **0.04** |

**Supplemental Table S3.** Results of a GLMM testing for an influence of condition (separation vs test trial) and identity of compartment (designated empty or test compartment) on the likelihood that pigs open a door. Significant predictors are indicated in bold.

| Predictor | Estimate | Std. Error | lower CI | upper CI | $\chi^2$ | df | *P* |
|---|---|---|---|---|---|---|---|
| (Intercept) | -0.434 | 0.473 | -1.361 | 0.494 | | | (†) |
| Condition_Test | 1.572 | 0.369 | 0.849 | 2.295 | | | (†) |
| Compartment_Test | -0.282 | 0.373 | -1.012 | 0.449 | | | (†) |
| Location compartment_Side§ | -0.232 | 0.278 | -0.777 | 0.313 | 0.689 | 1 | 0.405 |
| Test day_2¥ | 0.103 | 0.43 | -0.739 | 0.946 | 2.241 | 4 | 0.692 |
| Test day_3¥ | -0.179 | 0.422 | -1.007 | 0.648 | | | (Þ) |
| Test day_4¥ | 0.426 | 0.432 | -0.421 | 1.273 | | | (Þ) |
| Test day_5¥ | -0.085 | 0.461 | -0.988 | 0.818 | | | (Þ) |
| Daily test order in group_2nd# | -0.509 | 0.28 | -1.058 | 0.04 | 3.306 | 1 | 0.069 |
| **Condition_Test : Compartment_Test** | 1.496 | 0.576 | 0.367 | 2.624 | **6.956** | **1** | **0.008** |

† Not indicated due to limited interpretability (of intercept, or when there is a significant interaction)
Þ Refer to chi-sq statistic and p-value for 'Test day_2' for overall effect of predictor
§,¥,# Estimates refer to comparison with reference categories:
§ Location compartment_Front
¥ Test day 1
### Daily test order in group- 1st

**Supplemental Table S4.** Results of a LMM testing for a relationship between behavioural responses and changes in cortisol in trapped pigs. To test for effects on changes in cortisol, we interacted hormone sample context (pre- or post-trapped) with the predictors. Significant Interactions are indicated in bold.

| | Estimate | Std. Error | lower CI | upper CI | t or F statistic | df | P value |
|---|---|---|---|---|---|---|---|
| (Intercept) | 2.778 | 0.123 | | | | | † |
| Context_Before | -0.541 | 0.086 | -0.711 | -0.37 | | | † |
| Rate of distress signals‡ | 0.1 | 0.057 | -0.026 | 0.22 | | | † |
| Rate of window investigations‡ | 0.056 | 0.068 | -0.078 | 0.193 | | | † |
| Duration separation | -0.009 | 0.058 | -0.124 | 0.105 | | | † |
| Duration trapped¤ | -0.045 | 0.073 | -0.191 | 0.099 | | | † |
| Sex_Female§ | 0.014 | 0.084 | -0.154 | 0.182 | 0.164 | 60.3 | 0.871 |
| Test day 2¥ | -0.16 | 0.132 | -0.424 | 0.102 | 0.654 | 4, 61.96 | 0.626 |
| Test day 3¥ | -0.19 | 0.125 | -0.443 | 0.059 | | | (Þ) |
| Test day 4¥ | -0.138 | 0.139 | -0.42 | 0.138 | | | (Þ) |
| Test day 5¥ | -0.119 | 0.14 | -0.404 | 0.161 | | | (Þ) |
| Daily test order in group _2nd# | -0.038 | 0.093 | -0.222 | 0.147 | -0.409 | 64.47 | 0.684 |
| Time pig placed in compartment | 0.005 | 0.055 | -0.106 | 0.113 | 0.085 | 84.21 | 0.932 |
| **Context:Rate of distress signals** | -0.124 | 0.061 | -0.245 | -0.003 | **-2.041** | **70.05** | **0.045** |
| Context:Rate of window investigations | -0.097 | 0.075 | -0.246 | 0.053 | -1.284 | 69.82 | 0.203 |
| Context:Duration separation | -0.069 | 0.06 | -0.188 | 0.05 | -1.158 | 69.92 | 0.251 |
| Context:Duration trapped | 0.056 | 0.077 | -0.096 | 0.209 | 0.732 | 69.99 | 0.467 |

† Not indicated due to limited interpretability (of intercept, or because we are interested in evaluating interaction terms)

Þ Refer to chi-sq statistic and p-value for 'Test day_2' for overall effect of predictor

§,¥,# Estimates refer to comparison with reference categories:

§ Sex_Male

¥ Test day 1

### Daily test order in group-1st

‡ Data were square-root transformed

¤ Data were log transformed

**Supplemental Figure S1.** Diagram of the experimental set-up, showing two identical compartments that were attached to opposite sides of the home pen daily during familiarization and testing. The door of each compartment opens into the home pen when the handle is lifted high enough to release an inner latch.

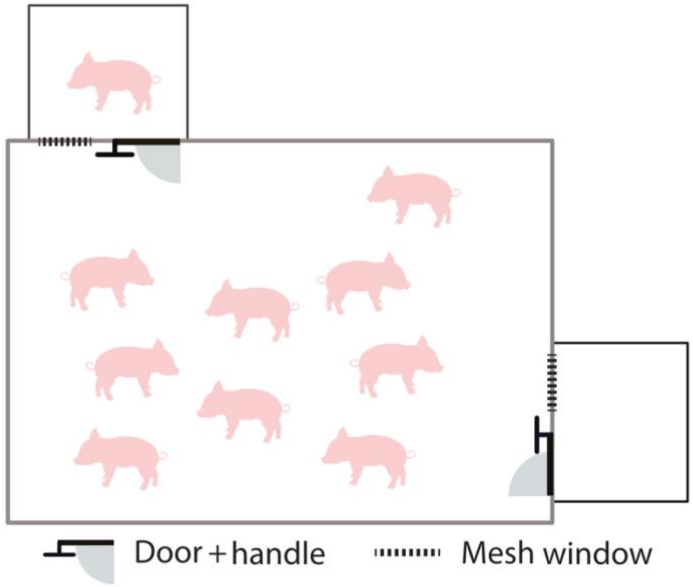

**Supplemental Figure S2.** View of a test compartment from inside the pen, including the door with handle and mesh window.

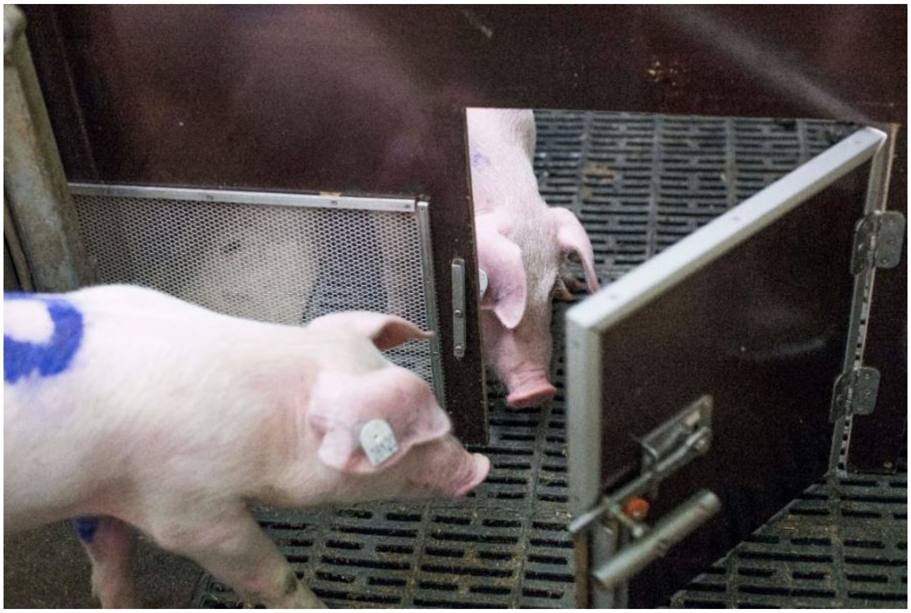

**Supplemental Figure S3.** Daily latencies (with medians and 25-75% interquartile ranges) for pigs to open compartments across familiarization days. Estimated marginal means and SEs for latencies to open the side compartment across days are indicated, with results of pair-wise post-hoc comparisons (Tukey's HSD) visualized using a compact letter display. Different letters indicate significant differences in latencies to open the side compartment. There were no differences in latencies to open the front compartment across days. For full model results see Table S1.

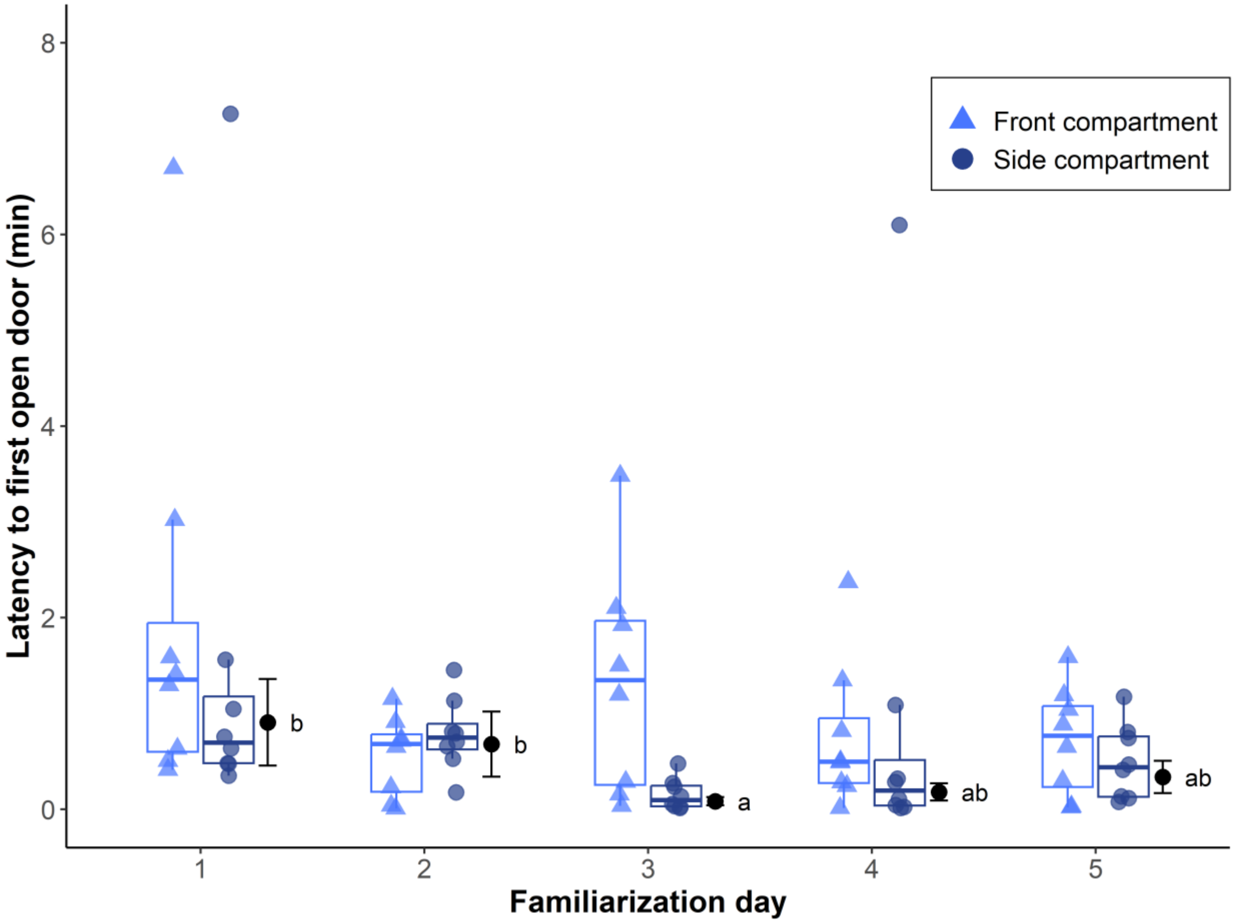

**Supplemental Figure S4.** Daily (mean and SD) proportion of pigs within each group who opened doors per familiarization day, and cumulative proportion of pigs to open a door at least once.

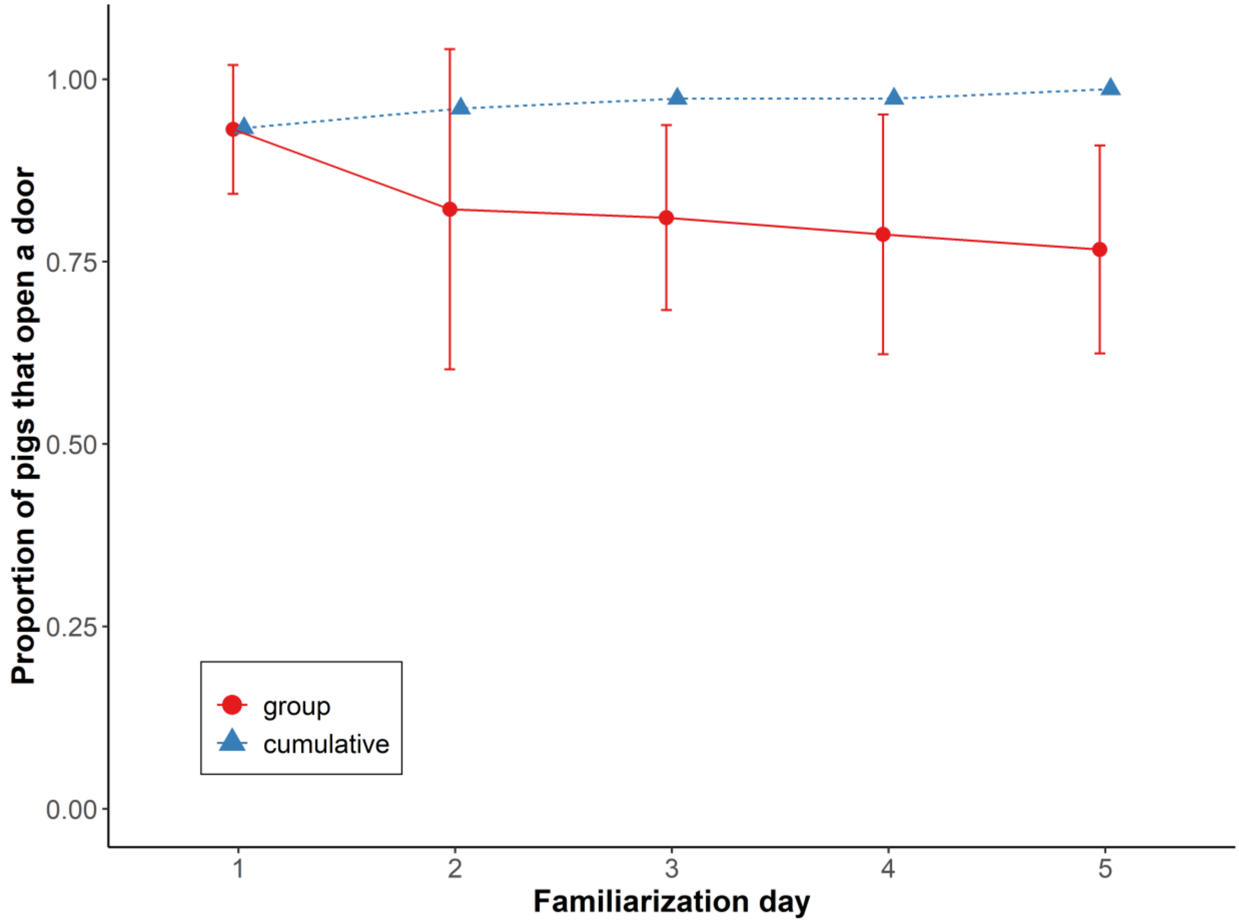

**Supplemental Figure S5.** Likelihood (medians and 25-75% interquartile ranges) for pigs to open compartments during test trials and during the preceding separation, when both compartments were empty. Points indicate likelihoods for each of n = 8 groups across their trials. Estimated marginal means and SEs are also indicated, with results of pair-wise post-hoc comparisons (Tukey's HSD) visualized using a compact letter display. Different letters indicate significant differences. For full model results see Table S3.

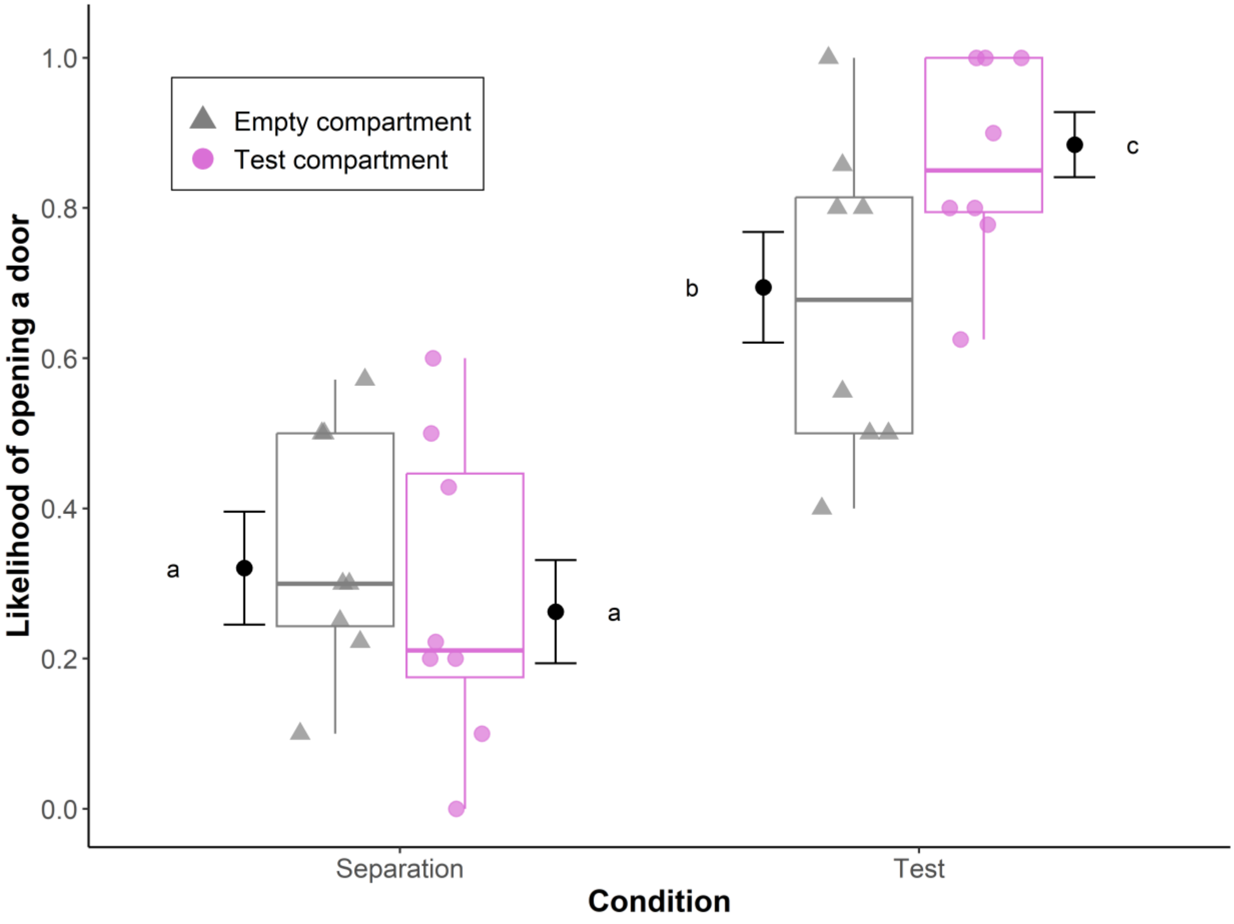

**Supplemental Figure S6.** Latencies (median and interquartile range) for helper pigs (indicated in circles) to release trapped pigs across test trials within each group. Colours indicate different social groups. Triangles indicate the trials where pigs were released by humans (not included in latency calculations).

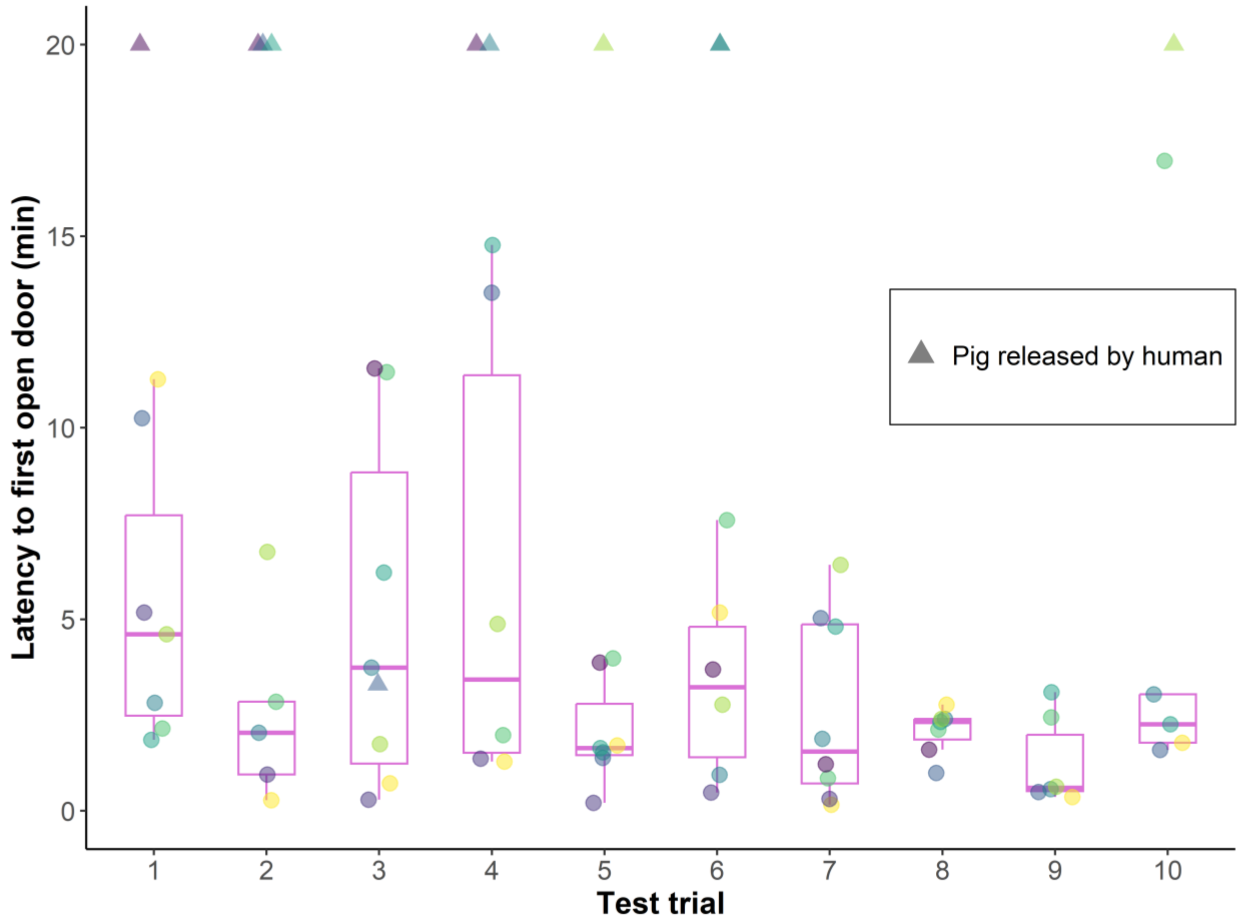

**Supplemental Figure S7.** Comparison of post-release salivary cortisol concentrations in trapped pigs and their helpers.

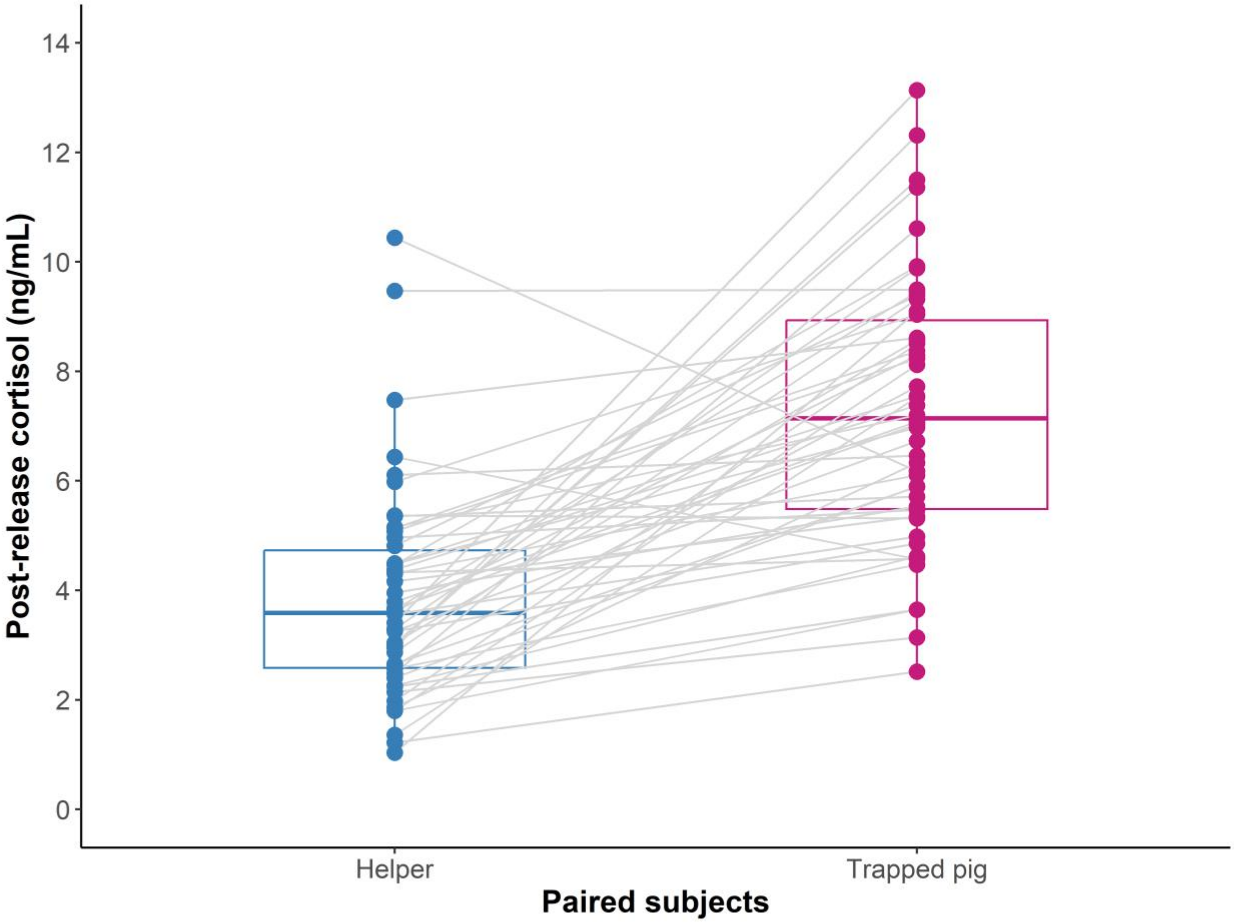

**Supplemental Figure S8.** Changes in salivary cortisol concentrations in pigs who exhibited fewer (≤ 1 per minute) or more (> 1 per minute) distress signals while trapped. Boxplots indicate medians and interquartile ranges. Rates of distress signals were analysed as a continuous predictor in the model. The categorization is included here to illustrate the results of the interaction between sample context and rates of distress signals. For full model results see Table S4.

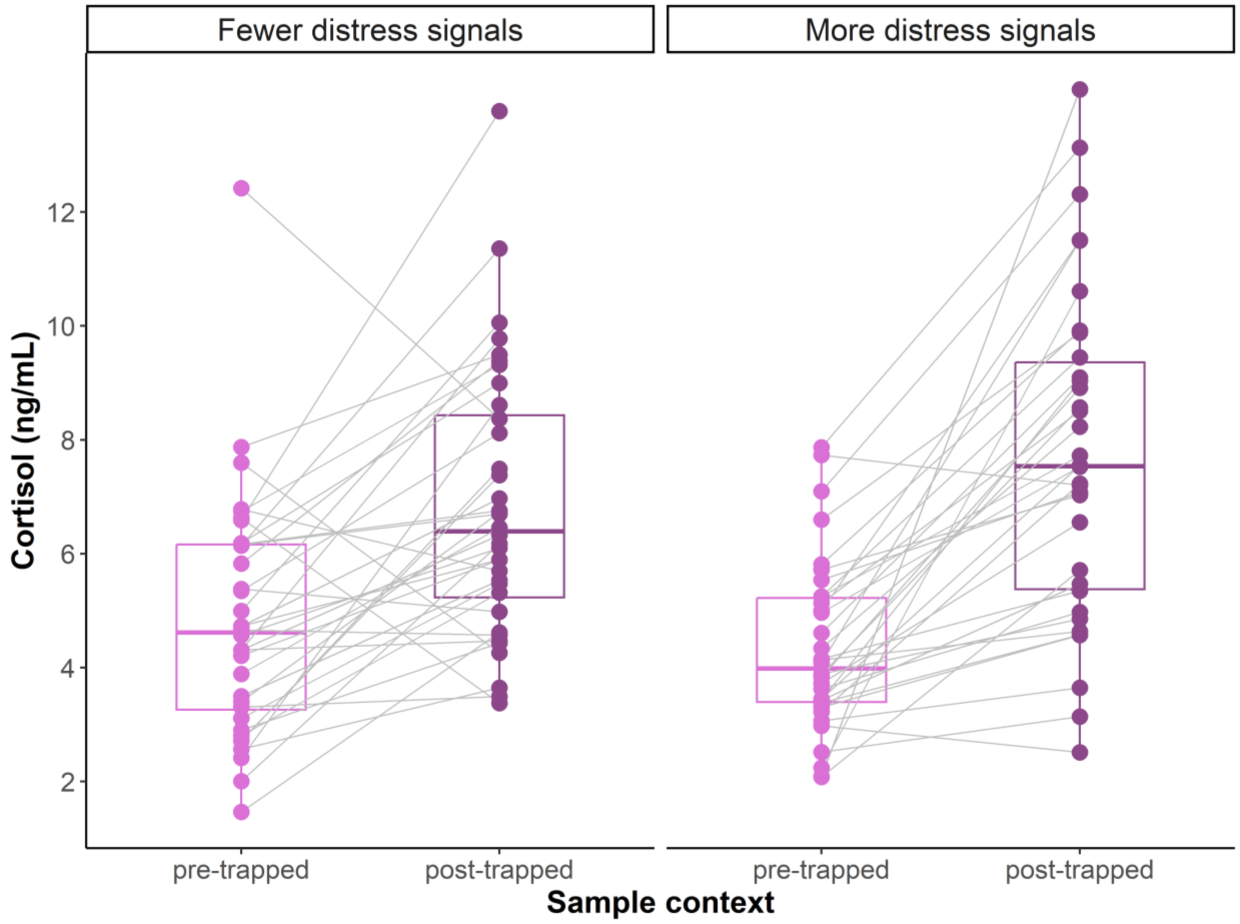

**Supplementary Video S1.** Top view of the test compartment, showing the behaviour of pig 4 in group 3B, shortly before being released.

https://lizamoscovice-research.com/proximate-factors-mediating-helping-behaviour-in-pigs/#v1

**Supplementary Video S2.** View of the test compartment from inside group 3B's home pen shortly before pig 4 was helped by pig 6. The control compartment is also in view to the left of the screen. Note that several potential helpers in the home pen display social nosing behaviour. Social nosing can occur during play and in social greeting contexts and is not indicative of aggression.

https://lizamoscovice-research.com/proximate-factors-mediating-helping-behaviour-in-pigs/#v2

**Supplementary Video S3.** Top view of the test compartment, showing the behaviour of pig 9 in group 2B, shortly before being released.

https://lizamoscovice-research.com/proximate-factors-mediating-helping-behaviour-in-pigs/#v3

**Supplementary Video S4.** View of the test compartment from inside group 2B's home pen shortly before pig 9 was helped by pig 8. The control compartment is also in view to the left of the screen.

https://lizamoscovice-research.com/proximate-factors-mediating-helping-behaviour-in-pigs/#v4
